# Rapid and high yield isolation of extracellular vesicles with purity by application of size exclusion fast performance liquid chromatography

**DOI:** 10.1101/2023.10.12.561425

**Authors:** Kshipra S. Kapoor, Kristen Harris, Kent Arian, Lihua Ma, Kaira A. Church, Raghu Kalluri

## Abstract

Extracellular Vesicles (EVs) have emerged as potential biomarkers for diagnosing a range of diseases without invasive procedures. Extracellular vesicles also offer an advantage compared to synthetic vesicles, for delivery of various drugs. However, limitations in segregating EVs from soluble proteins have led to inconsistent EV retrieval rates with low levels of purity. Here, we report a new high-yield (>95%) and rapid (<20 min) EV isolation method called <u>S</u>ize <u>E</u>xclusion – <u>F</u>ast <u>P</u>erformance <u>L</u>iquid <u>C</u>hromatography (SE-FPLC). We show SE-FPLC can effectively isolate EVs from multiple sources including EVs derived from human and mouse cells and serum. The results indicate that SE-FPLC can successfully remove highly abundant protein contaminants such as albumin and lipoprotein complexes, which can represent a major hurdle in large scale isolation of EVs for clinical translation. Additionally, the high-yield nature of SE- FPLC allows for easy industrial upscaling of extracellular vesicles production for various clinical utilities. Moreover, SE-FPLC enables analysis of very small volumes of blood for use in point-of-care diagnostics in the clinic. Collectively, SE-FPLC offers many advantages over current EV isolation methods and offers rapid clinical utility potential.

## Introduction

Advanced Therapeutic Medicinal Products (ATMPs) provide innovative approaches for designing solutions for diagnosing and combating complex diseases^1,2^. Promising breakthroughs in cutting-edge technologies, such as chimeric antigen receptor (CAR)-T cells^3–5^ and mRNA vaccines^6–9^, illustrate the potential of such unconventional thinking. These success stories demonstrate the utility of ATMPs and the need to explore novel therapeutic solutions for unmet medical needs. Extracellular vesicles (EVs) contain molecular cargo such as: proteins, nucleic acids, lipids, and metabolites reflecting their cell of origin^10–12^. EV membranes harbor lipid rafts characterized by high concentrations of cholesterol, sphingomyelin and ceramide, rendering them highly stable in body fluids and hence making them appealing for therapeutic purposes^13^. EVs also contain major histocompatibility complexes and can engage with the immune system in various antigen- specific ways^14^. Considering these benefits, EVs offer a promising platform for therapeutic development and in many cases, they can be altered to enhance their therapeutic potential thereby making them a versatile tool with broad applicability^15–18^. EVs have also garnered attention as potential disease biomarkers in liquid biopsy for cancer^19^ and neurological diseases^20^, among others. As EVs carry a diverse range of molecular cargo, they can serve as important biosignatures for disease diagnosis and therapeutic intervention. The potential diagnostic^21^ and therapeutic application^22^ of EVs highlights the importance for developing new protocols for their rapid isolation with purity to enable continued laboratory investigations and development as a valuable tool in disease management. However, the production of EVs, especially at an industrial scale, still represents a significant challenge, including product definition, low purity, and low yields^23–25^. Additionally, the lack of standardized EV isolation methods and the limited scalability of existing protocols for preparative purposes beyond the laboratory scale further compound these challenges. Therefore, our objective was to develop a scalable and reproducible method that meets the demands of the pharmaceutical industry to produce pure EVs.

Recent literature survey reveals that there are around 190 different methods reported for isolating extracellular vesicles (EVs) and over 1,000 unique protocols for extracting EVs from various sources of origin^26^. Differential ultracentrifugation (dUC)^27^ continues to be the most frequently employed method for isolating extracellular vesicles (EVs); however, this method necessitates lengthy protocols, limited by ultracentrifuge tube capacity in case of bulk isolation and often involves a trade-off between EV yield and purity^27–30^. Other approaches^31–36^ to EV isolation, such as polymer precipitation and immunoaffinity capture, have been explored to address the limitations of traditional methods, offering simplified procedures and shorter isolation times. However, polymer precipitation is prone to co-isolation of contaminating proteins, which can interfere with downstream analysis. Likewise, immunoaffinity capture methods may only isolate a subset of EVs, which introduces bias in the subsequent analysis^37^. While an alternative approach that avoids this potential limitation is utilizing bind-elute affinity chromatography for EV separation^38^, such an approach requires altering solvent conditions that can result in biological and chemical instability of biologics. These limitations underscore the need for a rigorous exploration of alternative EV isolation methods to ensure rapid and optimal results^39,40^.

This study presents a new ultrafast purification technique for EV isolation that overcomes the limitations of other methods. To confirm the efficacy of the SE-FPLC, we performed measurements on the isolated EV samples and address different published guidelines (MISEV2018)^41^ in the EV field. We thoroughly characterized the obtained EV fractions for enrichment of EV-specific markers and absence of non-EV proteins. To verify the EV purity standard protein detection methods were employed and appropriate size distribution was analyzed by nanoparticle tracking analysis (NTA), EV morphology was imaged by cryo-EM imaging. We compared the SE-FPLC performance with dUC, density gradient and size exclusion chromatography, and overall, SE-FPLC exhibited enhanced speed, yield and purity. We show that SE-FPLC delivers superior results with reduced processing time thereby aiding better economics and a lower carbon footprint.

## Results

### Development and characterization of SE-FPLC for high purity EV isolation

To achieve fast, efficient and pure EV isolation, we developed a SE-FPLC methodology that separates EVs based on size (**Figure 1A-E**). We investigated the EV purification by SE-FPLC utilizing AKTA pure 25 chromatography system (Cytiva). A FPLC compatible column qEV 10 IZON was attached to the cytiva system and we found that performing 0.22-micron pore size filtered distilled water wash and equilibrating the column with at least 5 column volumes of 0.22-micron pore size filtered phosphate-buffered saline (PBS) increased EV yield. One explanation for this outcome is that in-column washes are more effective at eliminating residual 0.1% sodium azide and ethanol which are present in the resin used for the column storage and preparation. Next, to characterize the separation efficiency, elution fractions from each peak in the Ultraviolet-visible absorption spectra at wavelength 280nm (**Figure 2A, 2D, 2G, 2J**) were continuously collected and tested for presence of EV- and non-EV- markers by western blot analysis. CD81, CD9 and Syntenin-1^42^ were used as EV markers and the glycolytic enzyme GAPDH and DNA binding protein Histone H3 were employed as non-EV markers respectively. We isolated EVs from 6 different cell lines representing different origins: bone marrow mesenchymal stem/stromal cells (MSCs); non-malignant epithelial (HEK293T); normal fibroblast (BJ); malignant epithelial cancer cells (Panc1, T3M4, KPC-689). Using trypan blue exclusion assay that allows for direct identification and enumeration of live (unstained) and dead (blue) cells in a given population, we observed low cellular death in the cells post serum starvation prior to EV isolation (**Supplementary Figure S1F**).

**Figure. 1.**
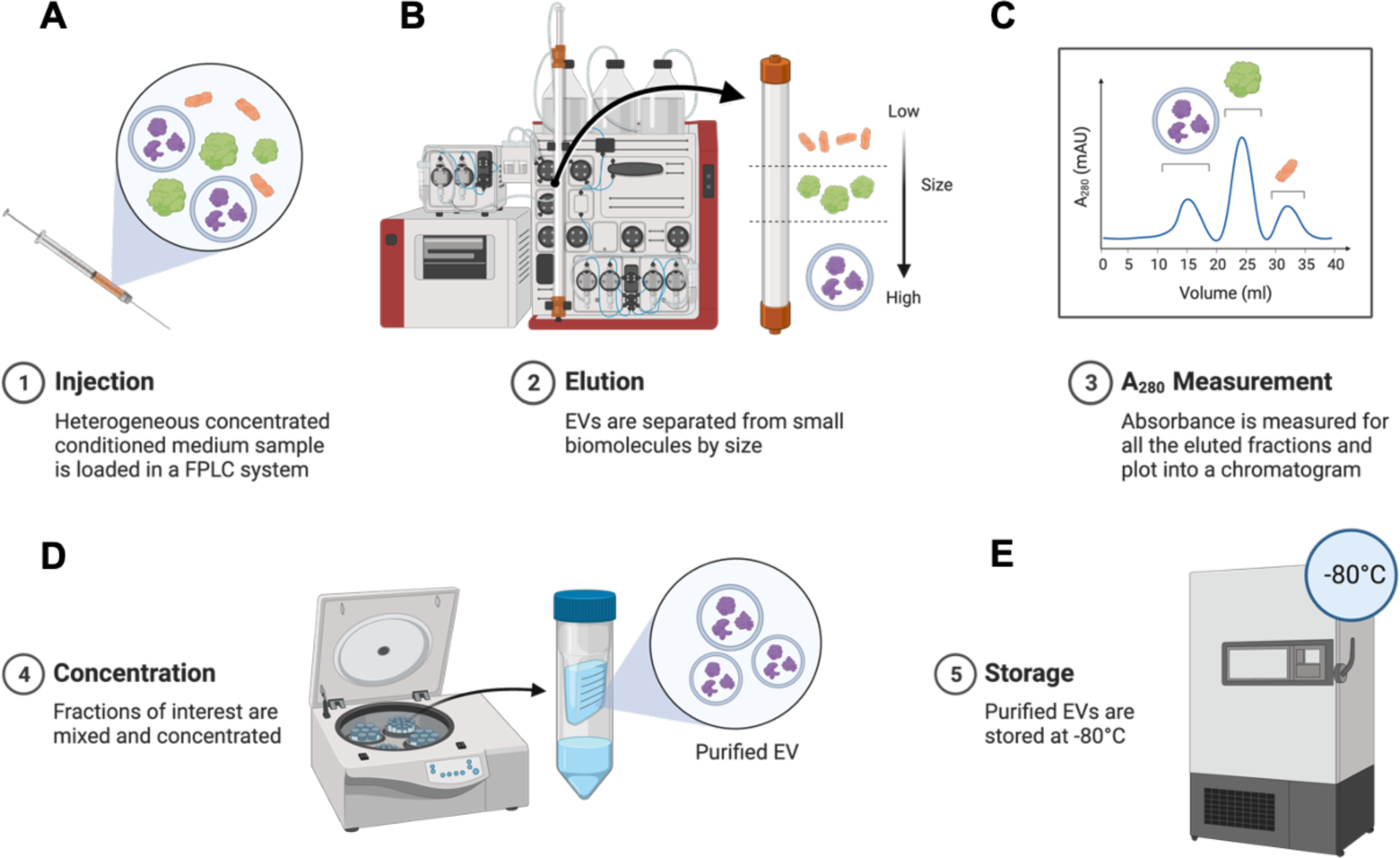
The size exclusion - fast protein liquid chromatography EV isolation system: SE-FPLC. (A-E) Schematic overview of the workflow. **(A)** A heterogeneous concentrated conditioned media sample mixture is loaded onto a FPLC compatible syringe. **(B)** After loading the concentrated conditioned media onto the SE-FPLC column EVs are separated from small biomolecules using the AKTA Pure chromatography system. **(C)** A built-in UV monitor is used to perform A_280nm_ measurements. The eluting fractions corresponding to each peak are collected via an automated fraction collector.**(D)** Fractions of interest are subsequently concentrated using a concentration filter. **(E)**Purified EVs are stored at -80°C until further usage and downstream analysis.

**Figure. 2.**
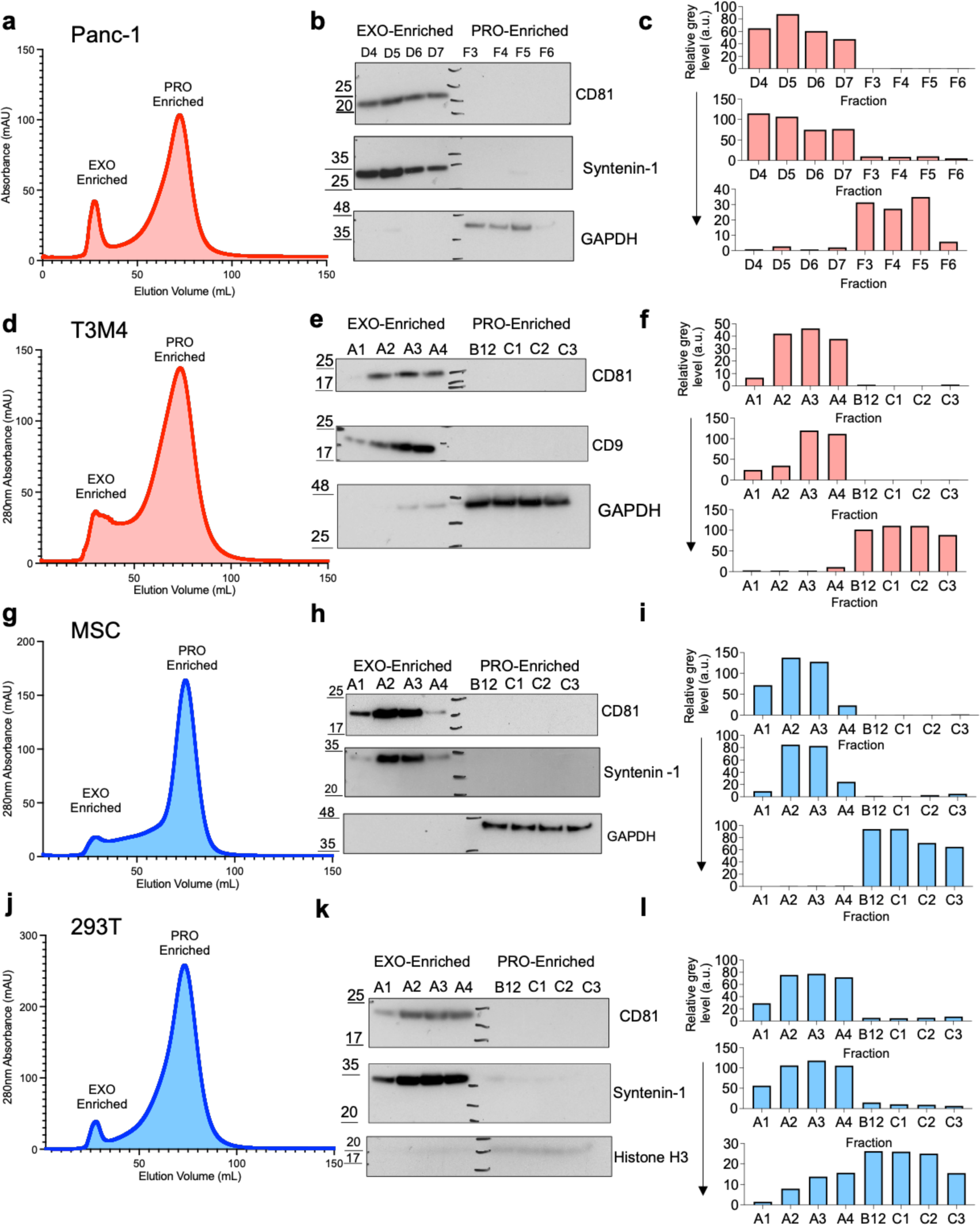
Biological characterization of SE-FPLC performance. UV-Vis chromatograph for EV isolation from **(A)** Panc1, **(D)** T3M4, **(G)** MSC, and **(J)** 293T conditioned media. Relative purity characterized by western blot results. Enrichment of exosomal proteins (CD81, CD9 and Syntenin-1) in EXO-enriched fractions indicating presence of the canonical small EV marker in these fractions. GAPDH and Histone H3, was used as a non-EV associated protein for verifying the EV purity for **(B)** Panc1, **(E)** T3M4, **(H)** MSC and **(K)** 293T respectively. Quantification of band intensities for respective fractions is shown in the right panel for **(C)** Panc1, **(F)** T3M4, **(I)** MSC and **(L)** 293T respectively. A low intensity of non-EV associated proteins in EV-enriched fractions indicates a higher EV purity.

Equal-sample-volume analysis (45μL) were used to characterize fractions that were EV-enriched for the presence of EV markers (**Figure 2B, 2E, 2H, 2K**). In different cell line isolated EVs, prominent band intensity for exosomal proteins corresponded to fractions in the first peak of elution. Further, the non-EV markers were absent in the fractions corresponding to the first peak of elution. GAPDH and Histone H3 were significantly enriched in fractions corresponding to the second peak of elution (**Figure 2B, 2E, 2H, 2K**) thereby indicating superior performance for separation of EVs from non-EV associated proteins.

The main performance metrics considered in this study were the processing time, EV purity and total yield. While most of the current protocols require at least 8 hours to isolate the samples, the SE-FPLC approach can isolate EVs in as little as 18 mins and in total 1 hour including the filtration steps common to all other isolation approaches (**Figure 3A**). Coomassie and silver staining of proteins further confirmed that SE-FPLC removed most protein contaminants across all model systems (**Figure 3B-3E** and **Supplementary Figure S6A-S6B**). Furthermore, the total EVs amount isolated via the gold standard techniques such as differential ultracentrifugation (dUC), density gradient (DG) and small- scale size exclusion chromatography (SEC) results in lower yield compared this novel SE-FPLC isolation method. A comparison between the yield (7-fold higher) and processing time (18 mins vs. 440 mins) for SE-FPLC versus dUC, DG and SEC are shown in (**Supplementary Figure S4A-S4B, Supplementary Figure S9A**) for comparison. In addition, we also compared the yield and processing time for commonly employed EV isolation techniques such as DG and SEC (**Supplementary Figure S5A-S5B, Supplementary Figure S9A**). The EVs isolated by SE-FPLC, dUC, SEC and DG were further inspected by cryogenic electron microscopy (Cryo EM) to determine EV morphology (**Figure 3G** and **Supplementary Figure S5F-S5I**). EVs isolated by various isolation methods displayed a similar size distribution between 40 to 200 nm (**Supplementary Figure S4C-4E** and **Supplementary Figure S5C-5D**).

**Figure. 3.**
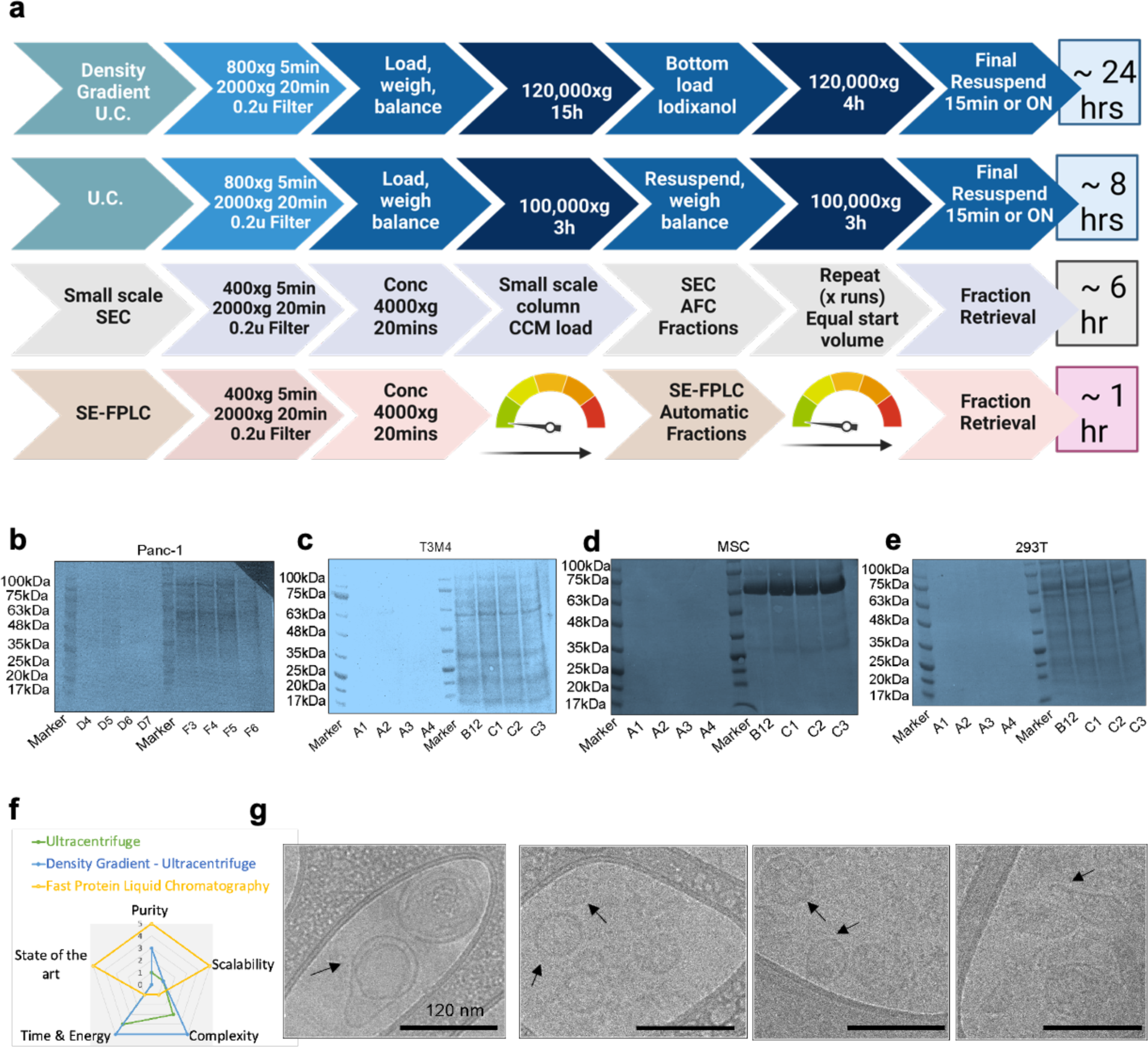
Characterization of SE-FPLC performance and processing time comparison to other methods of EV isolation. (A) Comparison between density gradient (DG), differential ultracentrifugation (dUC), small scale size exclusion chromatography (SEC) and size-exclusion fast protein liquid chromatography (SE-FPLC) detailed procedures and sample processing times. Equal-sample-volume analysis (45μL) of total proteins in each fraction by Coomassie staining for **(B)** Panc1, **(C)** T3M4, **(D)** MSCand **(E)** 293T EVs isolated by SE-FPLC. **(F)** Two-dimensional comparison of scalability, time and energy, purity and state-of-the art of SE-FPLC and other EV isolation methods.(G) Cryogenic electron microscopic images of EVs isolated from mesenchymal stem cells using SE-FPLC. Scale bars, 120 nm.

### SE-FPLC for rapid EV isolation exhibits scalability and high reproducibility

Next to evaluate the reproducibility and scalability of SE-FPLC, we processed independent biological replicates of Panc1, T3M4 and MSC EVs. The UV-Vis absorbance spectra indicated good technical reproducibility of SE-FPLC (**Supplementary Figure S1A-S1C**). We also determined the recovery rate of our SE-FPLC platform (**Supplementary Figure S4F-S4H**) and observed that approximately 98% of non-EV associated proteins were eliminated after SE-FPLC isolation of EVs and 88.47% of EVs were recovered in the first peak of elution. We isolated EVs from SE-FPLC and measured the protein amounts (μg/mL) concentration in EV-enriched peak fractions (**Supplementary Figure S10A**) and protein-enriched peak fractions (**Supplementary Figure S10B)**. The coefficient of variation of the measurements for the ratio of EV-enriched peak fractions and Pro-enriched peak fractions was 16.4%, thereby indicating reliable technical reproducibility of SE-FPLC.

### SE-FPLC method facilitates albumin depletion and removal of lipoproteins

Enriching extracellular vesicles (EVs) while effectively excluding highly abundant free proteins such as albumin and lipoproteins of similar density, such as high-density lipoprotein particles (HDLs), poses a significant technical challenge. Given the relatively low abundance of EVs compared to free proteins and lipoproteins, achieving their enrichment while avoiding co-isolation of these impurities remains challenging^43,44^.

SE-FPLC was able to enrich EVs from both free proteins (albumin – the most abundant free protein) and lipoprotein (HDL) with the ability to quantify the separation efficacy. Our SE-FPLC approach aided in albumin depletion in as little as 18 mins (**Figure 4A**). Whereas commercially available albumin-depletion kits took over 1 hour and 30 minutes for the removal of albumin from serum (**Figure 4B**). The separation of contaminating albumin by SE-FPLC largely enhances downstream analysis, including western blotting, and enables the detection of specific proteins as potential biomarkers and/or therapeutic targets. To determine if the albumin depletion was successful, we quantified albumin levels in both EV-enriched and non-EV-enriched fractions across multiple model systems (**Figure 4C-4E**). Our approach greatly improved the albumin- depletion process for purification of EVs, thereby improving various downstream analysis.

**Figure. 4.**
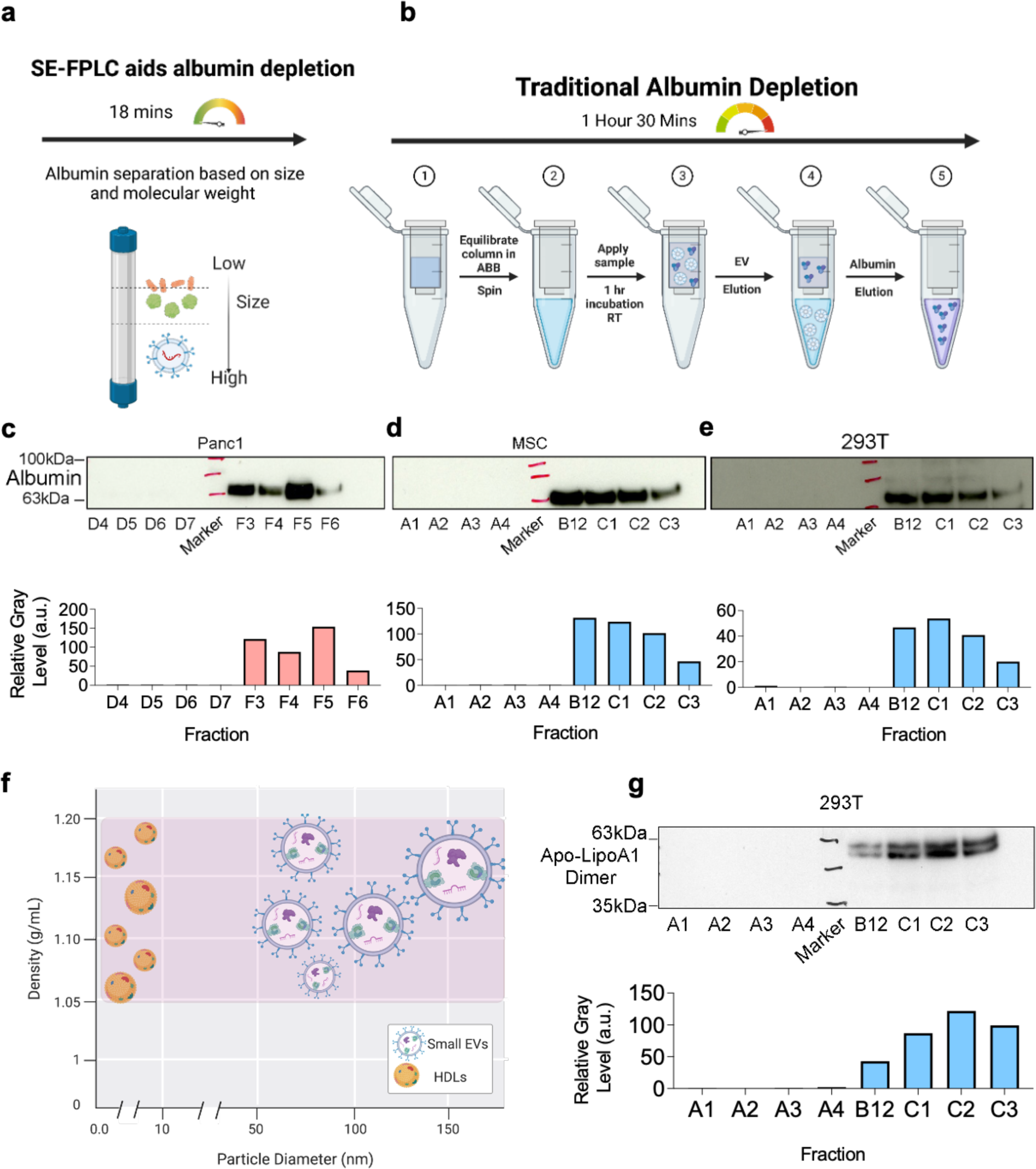
SE-FPLC method facilitates albumin depletion and removal of lipoprotein complexes. (A) Schematic of albumin depletion in one-step and processing time via the SE-FPLC method. **(B)** Schematic of traditional albumin depletion procedure and thetraditional protocol processing time. Extracellular vesicle samples were isolated by SE- FPLC approach and equal-sample volumes (45μL) of each fraction were analyzed by western blot analysis for presence of albumin in **(C)** Panc1, **(D)** MSC and **(E)** 293T derived EVs. The relative intensity quantification of the western blot band was analyzed by ImageJ and is shown below each respective blot. **(F)** Comparison of density (g/mL) and particle diameter (nm) of high-density lipoprotein (HDL) particles and small extracellular vesicles (sEVs). **(G)** Western blot analysis of EVs isolated from 293T cells via SE-FPLC approach. Apo-LipoA1 (marker for HDL particles) are enriched in the non-vesicular enriched fractions.

EVs such as exosomes have been reported to have similar density as small HDL particles both belonging in the range of 1.05 – 1.21 g/mL (**Figure 4F**). Thus, due to the similar densities there are high chances of co-isolating both small EV and HDL particle populations using traditional approaches such as density gradient ultracentrifugation^43,45–47^. SE-FPLC approach demonstrated the ability to separate EVs and HDL particles by size, HDL particles (5 – 10 nm in diameter) and sEVs (50 – 220 nm) were easily fractioned in one-step due to size differences via the SE-FPLC. Apolipoprotein A1 (Apo-Lipo A1) a marker for HDL particles was used to quantify the efficacy of separation between EV and HDL particles. Dimerized apo-lipo A1 was enriched in the non-EV fractions (**Figure 4G**) hence confirming the SE-FPLC partitions HDL particles in the non-EV enriched fractions away from the purified small EV population.

### Isolating serum EVs via SE-FPLC

To determine the versatility of SE-FPLC, we evaluated the ability to isolate and purify EVs from human serum. Serum-derived EVs can function as liquid biopsies for monitoring disease-associated changes as they are released into the bloodstream from their cells of origin in the tissue^48^. We isolated EVs from healthy serum samples using SE-FPLC. Age and sex data for samples employed in this study are reported in Supplementary Table 1. The UV-Vis absorbance spectra of healthy donor samples are represented in **Figure 5A**. Due to high-yield and high-purity, SE-FPLC reduced biofluid starting volume requirements to 500 μL.

**Figure. 5.**
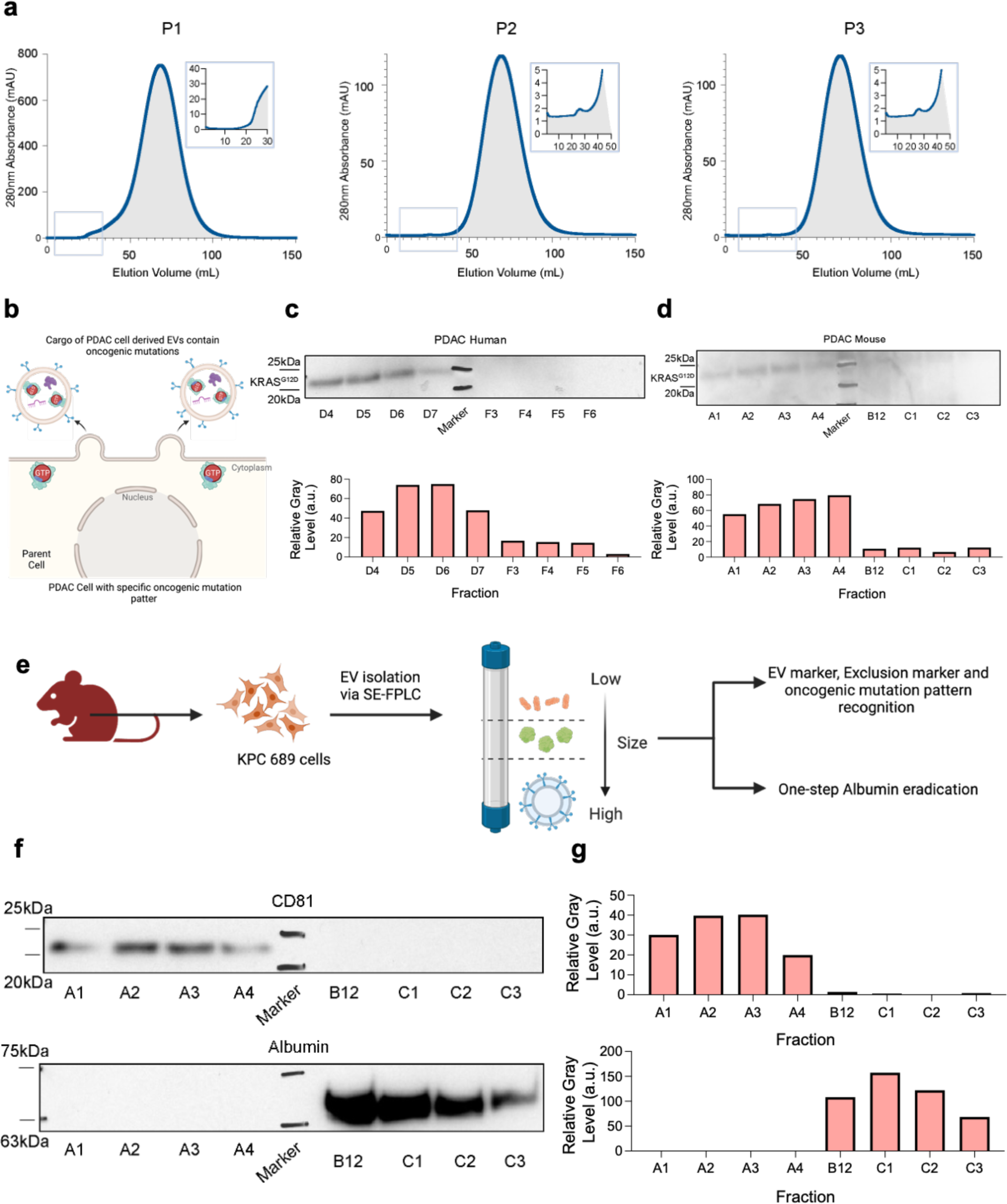
Validating versatility of SE-FPLC method and detection of mutant KRAS in SE-FPLC isolated EVs. (A) UV-Vis chromatographs for EV isolation from serum of n=3 healthy individual donors. **(B)** Schematic of compartmentalization of mutant KRAS in the PDAC cell derived EVs. **(C, D)** Immunoblot analysis of EVs isolated from PDAC human (Panc1) and PDAC mouse (KPC689) cells for KRAS^G12D^ protein. KRAS^G12D^ was associated with EV enriched fractions. Quantification of relative intensity of the bands was analyzed via ImageJ and is shown below the respective blot. **(E)** Workflow of EVs isolated from KPC689 mouse cell lines. **(F)** Western blot analysis to study the purity of mouse cell line derived EVs. Equal-sample-volume analysis (45μL) to study the expression of common exosomal protein (CD81) and expression of most abundant free floating protein albumin. At equal-sample-volume analysis high intensities of CD81 in EV-enriched fractions indicated higher EV yield and low intensity of albumin in EV-enriched fractions indicates a higher EV purity. **(G)** Quantification of relative intensity of bands from the western blot was analyzed by ImageJ.

The serum EVs isolated by SE-FPLC were further inspected by cryogenic electron microscopy (Cryo – EM) (**Supplementary Figure S8A-S8F**). Majority of the serum isolated EVs revealed size integrity between 50 and 200 nm. About 70% of single-molecule independent EVs were in size range between 30 and 140 nm. About 25% of the single-molecule independent EVs ranged between 150 and 220 nm (**Supplementary Figure S8G**).

### SE-FPLC isolated EVs from pancreatic cancer cell lines detect mutant KRAS in EVs

To highlight the performance for practical cancer applications of SE-FPLC, we isolated EVs from pancreatic ductal adenocarcinomas (PDACs) cell lines using automated SE-FPLC. As KRAS mutation is associated with greater than 95% of PDAC patients^49^, we profiled SE-FPLC isolated EVs for mutated Kras (e.g., KRAS^G12D^) using mutant specific KRAS antibodies. We identified KRAS^G12D^ protein in EV-enriched fractions (**Figure 5B-5D**) across EVs isolated from both human and mouse PDAC cell lines. Additionally, we confirmed enrichment of EV associated markers CD81 and successful albumin depletion in EVs isolated from mouse PDAC cell line (**Figure 5E- 5G**).

We compared our SE-FPLC approach with SEC using small column volume and found the presence of oncoprotein KRAS^G12D^ in EV-enriched fractions (**Supplementary Figure S5E**). To validate the assay specificity, we isolated EVs from pancreatic cell lines with wild-type (WT)- KRAS (**Supplementary Figure S7C**). We observed no signal for oncoprotein KRAS^G12D^ in the EV-enriched fractions.

## Discussion

A quest for developing a reliable, scalable, rapid, and pure EV isolation method is a highly sought-after goal in the field to further enhance our understanding of the fundamental biological and translational significance of EVs in health and disease. Here we present SE-FPLC, a platform for fast, high performance EV purification^25,32,50^. The SE-FPLC method enables separation of EVs based on size while maintaining their natural morphology and protein marker characteristics. Moreover, the scalability of this platform makes it a feasible option for large-scale EV isolation from various types and volumes of biofluids from any species. Multiple studies have reported that various EV isolation methods have a tendency to co-purify different subpopulations of EVs due to their different principles^51^. Our western blot analysis revealed that the EVs isolated through SE-FPLC exhibited the highest signal intensities for specific exosomal proteins, while demonstrating the lowest signal intensity for selected non-EV associated proteins. These findings underscore the effectiveness of SE-FPLC in achieving efficient and pure isolation of EVs. Furthermore, the automatic operation and fraction collection feature of SE-FPLC guarantee a consistent workflow and reproducible results across various sample types, volumes, and EV concentrations. Thus, the integration of SE-FPLC with an automated system presents a convenient and robust approach for the rapid isolation of EVs with high purity and yield, surpassing current EV purification methods.

By employing SE-FPLC for purification of EVs from different cell line models and serum from patients we show the versatility of our new platform. By profiling EVs isolated from PDAC positive samples, we found enrichment of mutant oncoprotein – KRAS^G12D^ in EV enriched fractions, indicating the feasibility of SE-FPLC in clinical applications. We have identified certain features that have the potential to enhance the functionality of SE- FPLC in future studies. First, the current form of SE-FPLC is limited to single-column isolation. Multi-column approach can be implemented for generating EVs in a continuous fashion to reach highest productivity at an industrial scale. Second, bioreactor systems and hollow-fiber membranes can be utilized for concentrating and filtering cell culture conditioned media for the large-scale production of EVs. Third, the integration of SE- FPLC with downstream detection and analysis technologies such as automated capillary western blot, immunoassay, and quantitative PCR would be advantageous for fast and seamless enrichment and investigation of EVs. Additionally, it is essential to optimize the isolation workflows tailored to the characteristics of each specific sample type.

To summarize, we introduce the SE-FPLC methodology for efficient and rapid isolation of EVs with high purity and yield from both cell line models and biofluids, which has the potential to accelerate EV research in life sciences and facilitate translation to clinical applications. The scalability of our approach, enabled by the capability to process large sample volumes and the potential for automation, makes it attractive for industrial applications.

## Materials and Methods

### Cell culture

The cell lines employed in our study (HEK 293T, Panc1, BJ) were obtained from American Type Culture Collection (ATCC), (T3M4) was obtained from the cell bank at RIKEN BioResource. HEK 293T, Panc1, BJ Fibroblasts and T3M4 cell lines were validated by the Cytogenetics and Cell Authentication Core at MD Anderson. For HEK 293T and BJ the culture media was DMEM (Corning) supplemented with 10% fetal bovine serum (FBS) (Gemini) and 1% penicillin-streptomycin (Corning). For T3M4, Panc1, KPC-689 the culture media was RPMI 1640 (Corning) supplemented with 10% FBS and 1% penicillin-streptomycin. The KPC689 PDAC cancer cell line was isolated from an autochthonous pancreatic tumor of Pdx1^cre/+^; LSL-Kras^G12D/+^; LSL-Trp53^R172H/+^ (KPC) mice as described previously^52^. Bone marrow derived MSCs were obtained from Cell Therapy Laboratory at the University of Texas at MD Anderson Center. MSCs were cultured in alpha MEM (Corning #10-012-CV) supplemented with 2 U/mL heparin (Sigma H3149-100KU), 1% L-glutamine (Corning #25-005-CI), 1% Pen-Strep (Corning #30-002-CI), 1% non-essential amino acid (NEAA) (Corning #11140050), and 5% PLT Max (EMD Millipore #SCM141). All the cell lines were regularly tested for mycoplasma contamination using the LookOut Mycoplasma PCR detection kit (Product no. #MP0035 Sigma-Aldrich) and maintained in humidified cell culture incubators at 37°C and 5% CO_2_.

### EV production

Cultured cells at a confluency of about 80% were thoroughly washed with PBS and subjected to serum-free medium for 48h. The conditioned medium (CM) of the serum-starved cells was harvested and subjected to low g centrifugation steps: 400 x g for 10 mins to pellet cells. Supernatant was centrifuged at 2000 x g for 20 min to remove cellular debris and apoptotic bodies. Next, the CM was filtered using a 0.2 μm pore size filter flask (Fisher Scientific). After the initial processing, extracellular vesicles were isolated using different isolation methods.

### EV isolation

Four isolation techniques were employed for EV purification: Size Exclusion – Fast Protein Liquid Chromatography (SE-FPLC), OptiPrep – Density Gradient (DG), differential Ultracentrifugation (dUC) and Size Exclusion Chromatography (SEC).

### SE - FPLC

All separation and purification steps were performed on the most frequently utilized AKTA pure 25 chromatography system (Cytiva) equipped with the following components with cytiva numbers, injection valve (V9-inj), sample pump (S9) with 7 port inlet, Inlet A(V9-IA), and Inlet B (V9-IB). Active monitoring parts were multi-wavelength UV monitor (U9- M), conductivity meter (C9) and pH meter (V9-pH). The UV monitor measured absorbance in the UV/Vis range from 190 nm to 700nm. The 280nm wavelength was chosen to evaluate the purification performance and process. The flexible fraction collector (F9-C) that contains cassettes for 96 deep-well plate was applied for all the fractions. The purifications were facilitated by using UNICORN software version 7.3.

After filtration the CM was concentrated using a sterilized Amicon Ultra-15 10kDa filter following the manufacturer’s protocol. The FPLC-compatible column was attached to the FPLC system and then washed with 0.22-micron pore size membrane filtered 20% Ethanol and distilled water. Next the column was equilibrated with Buffer A (phosphate- buffered saline 1x) for 5 column volumes (CVs). In order to clean the sample loop, the procedure was the following to avoid sample going directly to the waste: 1) chose the system in “load”, wash the sample loop with PBS using a syringe with volume capacity larger than the loop. 5ml syringe was selected and left to the injection port. 2) Switch system to “inject”, detached the wash syringe and put the sample syringe in the injection port, avoid pushing the plunger. 3) Change the system to “load” and slowing push the plunger of sample syringe. 4) Switch the system back to “Inject”. The concentrated conditioned media were applied and loaded to the column. The column was washed with 1.5 CVs of 1X PBS. The peak fractionation (2 mL) was gathered by the 96 deep well plates using the F9-C fraction collector. EV elution peak and non-vesicular matter elution peak were further confirmed by measuring the absorbance of the fractions in real-time by a multiwavelength UV monitor (U9-M). The western blot analysis was carried out for vesicular and exclusion markers on the FPLC fractions and the number of particles and total protein in each fraction were determined. In short, equal volumes of collected fractions (45μL) were subjected to western blotting. The EV-enriched SE-FPLC fractions were further concentrated using the sterilized Amicon Ultra-10 100kDa filter and EV morphology and size were confirmed by high resolution cryogenic electron microscopy imaging. The FPLC running buffer (Buffer A) consisted of potassium dihydrogen phosphate (0.144 g/L), sodium chloride (9 g/L), disodium phosphate (0.795 g/L) at pH 7.3 to 7.5.

### Size Exclusion Chromatography (SEC)

After filtration the CM was concentrated using a sterilized Amicon Ultra-15 10kDa filter. 500 μl of concentrated CM (CCM) was overlaid on 70 nm qEV 500 size exclusion columns (Izon, SP1) for separation. A dedicated column was used for each cell line, and each column was used up to its limit. The column was flushed in between samples using filtered PBS and filtered 20% ethanol. For each EV sample, twenty-four fractions of 500 μl were collected using the Izon automated fraction collector (AFC v1). The fractions F7 to F25 were analyzed by western blot to probe for EV-enriched and non-EV enriched fractions. The EV-rich fractions were F7-F10 and EV-Poor fractions were F15-30. Pooled fractions (7-10) and (11-30) were then concentrated using 10 kDa cutoff filters (Amicon Ultra-15, Millipore). Each fraction was subjected to protein concentration measurement, Nanoparticle Tracking Analysis (NTA) and western blot analysis following manufacturer’s instructions. The EV-enriched fractions (F7-F10) were further concentrated using sterilized Amicon Ultra-15 10kDa filter analyzed by high-resolution cryogenic electron microscopy.

### Differential Ultracentrifugation (dUC)

For EVs isolation by dUC method, harvested conditioned media was subjected to ultracentrifugation at 100,000 x g for 3h at 4°C using a SW32 Ti Beckman Coulter rotor. After ultracentrifugation, the supernatant was discarded, and the EV pellet was resuspended in PBS. For performing cryogenic electron microscopic imaging EVs were spun down and washed in PBS by ultracentrifuge at 100,000 x g for 3h at 4°C. For performing the wash step a SW41 Ti rotor (Beckman Coulter) was employed. For MSC EVs isolation the CM was subjected to ultracentrifugation (Sorvall WX 100+ ultracentrifuge) at 100,000 x g for 3h at 4°C using ThermoFisher T-647.5

### Density gradient (DG)

EVs that were isolated by UC (1 ml) were further purified using a density gradient purification employing OptiPrep Medium from Sigma Aldrich. OptiPrep gradients were prepared using PBS with specific percentages and volumes for each gradient: 12% (2 ml), 18% (2.5 ml), 24% (2.5 ml), 30% (2.5 ml) and 36% (2.5 ml). The EVs were mixed with the OptiPrep medium to achieve a final concentration of 36% and were loaded at the bottom of a small (11mL) ultracentrifuge tube. The subsequent gradient fractions were gently layered on top in descending order: 30%, 24%, 18% and finally 12%. The gradients were then ultracentrifuged at 120,000 x g for 15 h using SW41 Ti rotor from Beckman Coulter. The following day, 12 fractions of 1 ml each were collected and labeled as F1 to F12. Each fraction was then washed in PBS with a 12-fold dilution and subjected to ultracentrifugation at 120,000 g for 4 h. Subsequently all the fractions were resuspended in PBS and the EV rich fractions (F1-F6) were utilized for Cryogenic electron microscopy imaging and NTA analysis.

### Clinical Samples

Serum samples from healthy participants were considered deidentified discarded material and exempt from requiring approval from the Institutional Review Board (IRB) of University of Texas at MD Anderson Center, and informed consent was obtained from all participants. Serum samples were collected from each participant, and relevant patient information can be found in Supplementary Table 1. Each serum sample was centrifuged at 400g for 10 mins followed by 2000g for 20 mins at 4°C. The resulting supernatant was then filtered through a 0.22 um pore size syringe filter followed by SE-FPLC EV isolation. Post SE-FPLC EV-enriched fractions were combined using a Amicon Ultra-15 100kDa cutoff concentration filter and serum EVs were subjected to high resolution cryogenic electron microscopy characterization.

### NTA

The concentration and the size distribution of the EVs was measured based on their brownian motion using a Nano Sight LM10 (Malvern), Blue 488 laser and a highly sensitive sCMOS camera. During measurements, temperature was set and kept constant at 25°C. For each acquisition, a 90 s delay followed by three captures of 30 s each was employed. The average values of three captures were used to determine the nanoparticle concentration and the mode and mean of the size distribution.

### Western Blot analysis

For western blotting EVs were loaded on 4-12% precast polyacrylamide mini gels (Invitrogen) for electrophoretic separation of proteins. Protein transfer was performed on methanol-activated polyvinylidene fluoride (PVDF) membranes by Trans-Blot Turbo Transfer system (1704150; Bio-Rad). After transfer the membranes were blocked in 5% BSA in TBS with 0.1% Tween-20 for 1 hour at room temperature. Post blocking the membrane was incubated in primary antibody overnight on a shaker. Next day, secondary antibodies were incubated for 1h at room temperature. All the primary and secondary antibodies and their concentrations used are listed in Table 2. Post primary and secondary antibody incubations the membranes were washed with TBS containing 0.1% Tween-20 on a shaker, three times at 10-min intervals. The visualization of the membrane was performed with West-Q Pico ECL solution (Gen-depot) following the manufacturer’s instructions. Amersham Hyper film (GE Healthcare) was used for capturing the chemiluminescent signals. All the original and full-length scans of blots are provided in Supplementary Data.

**Table 2:** Antibodies used for western blot analysis.

| Protein Target | Host species | Dilution | Vendor and Cat. No. |
| --- | --- | --- | --- |
| Syntenin - 1 | Rabbit | 1:2000 | Abcam, ab133267 |
| CD81 | Mouse | 1:1000 | Santa Cruz, sc-166029 |
| GAPDH - HRP<br>Conjugated | Rabbit | 1:1000 | Cell Signaling<br>Technology, CST -<br>#8884 |
| Histone H3 | Rabbit | 1:1000 | Abcam, ab201456 |
| Syntenin - 1 | Rabbit | 1:1000 | Abcam, ab19903 |
| CD9 | Rabbit | 1:1000 | Abcam, ab263019 |
| Albumin | Rabbit | 1:1000 | Cell Signaling<br>Technology, CST -<br>#4929 |
| Apolipoprotein A1 | Rabbit | 1:1000 | Abcam, ab52945 |
| Anti-mouse HRP-conjugated | - | 1:5000 | R&D, HAF007 |
| Anti-Rabbit HRP-conjugated | - | 1:5000 | Cell Signaling<br>Technology, CST -<br>#7074S |

### Cryogenic Transmission Electron Microscopy

EVs were washed and resuspended in 50 μl of 1x PBS with a minimum concentration of 5.5e10 EV/50 μl. Quanti-foil mesh grids and Lacey Carbon only (400 Mesh Cu) grids were glow discharged for 120 sec just before the vitrification step. During vitrification, approximately 3-5 μl of the EVs samples were applied on to the grids, blot time was optimized for 2-5 sec, blotting force was optimized between (1∼4) to obtain good-quality EV film and followed by snap freezing into liquid ethane using the FEI Vitrobot Mark IV system. The frozen grids were placed in a cryo specimen holder and then transferred to liquid nitrogen for the examination. The samples were imaged using a JEOL 2010 Cryo-TEM (200kV, LaB6 filament), with a Gatan 626 cooling holder operated at -180 °C. Mini Dose System (MDS) were used for image processing. The vitrification and imaging steps were performed at Shared Equipment Authority (SEA), Rice University. The DM4 images obtained were converted to TIF and further image analysis was performed. Custom Python codes were utilized for quantifying the segmented EVs for size distribution, length of major and minor axis and eccentricity of the particles.

### Statistics

Prism 9.2.0 software was used for graphical representation and statistical analysis. The error bars in the graphical data represents mean ± standard deviation as indicated in the figure legends. Normal distribution was assessed using the Shapiro-Wilk test and the datasets passed the normality test. For comparison of two groups statistical significance was determined using an unpaired t-test. Value of p<0.05 indicated statistical significance. The tests used to determine the statistical significance are indicated in the figure legends along with the exact p-values.

## Data availability

The main data supporting the results of this study are available within the manuscript and supplementary information files. The raw data files are available for research purposes from the corresponding author upon reasonable request. Source data are provided with this paper.

## Code availability

Custom Python scripts used for quantifying cryogenic electron microscopic images are available from corresponding author on request.

## Author contributions

Kshipra S. Kapoor – Conceived the idea, conceptualization, methodology, investigation, data analysis, prepared displays communicating datasets and writing.

Kristen Harris – investigation Kent Arian – investigation Lihua Ma – investigation

Kaira A. Church – investigation

Raghu Kalluri – Conceived the idea, conceptualization, project oversight and supervision, resources, funding, and writing.

## Acknowledgements

This work was supported by a gift from Fifth Generation, Inc. (“Love ‘Tito’s’”) and funds from Sid W. Richardson Foundation to RK. EV work in the Kalluri lab is supported by NIH R35CA263815, NIH P40OD024628 and NCI (R01CA231465-02). We are grateful to Dr. Wenhua Guo for training KSK on cryogenic electron microscopy. We are grateful for Dr. Kathleen M. McAndrews for the insightful discussions. LM is thankful to the Shared Equipment Authority at Rice University for the support. Graphical figures in this work were created using BioRender.

## Supplementary Figure

**Supplementary Figure. 1 | Reproducibility of EV isolations across multiple model systems using SE-FPLC.** UV chromatograms from independent EV isolations showing good peak separation between EVs and non-EV associated proteins. **(A)** n=3 independent biological replicates for Panc1 cell derived EVs. **(B)** n=2 independent biological replicates for T3M4 cell derived EVs. **(C)** n=4 independent biological replicates for MSC cell derived EVs. **(D, E)** UV-Vis chromatograms for EV isolated from 293T and BJ fibroblasts. SE-FPLC isolation shows consistency between batches. **(F)** Quantification of percent viability (trypan blue exclusion) in the parental cells upon serum-starvation for EV production. Violin plots represent n=18 samples for Panc1, T3M4 and MSC group and n=6 samples for 293T group. Short-dashed line represents the median of the datapoints in the Violin plots.

**Supplementary Figure. 2 | Full length unprocessed blot pertaining to** Figure 2**. (A)** Full length blots corresponding to Panc1 CD81, Syntenin-1 and GAPDH associated with main Figure 2b. **(B)** Full length blots corresponding to T3M4 CD81, CD9 and GAPDH associated with main Figure 2e. **(C)** Full length blots corresponding to MSC CD81, Syntenin-1 and GAPDH associated with main Figure 2h. **(D)** Full length blots corresponding to 293T Syntenin-1, CD81 and Histone H3 associated with main Figure 2k.

**Supplementary Figure. 3 | Full length unprocessed blot pertaining to** Figure 4 and**5. (A)** Full length blots corresponding to Panc1 Albumin, MSC Albumin and 293T Albumin associated with main Figure 4c, d and e respectively. **(B)** Full length blots corresponding to 293T Apolipo-protein A1 (HDL particle marker), Panc1 RAS^G12D^ and KPC689 RAS^G12D^associated with main Figure 4g, Figure 5b and 5c respectively. **(C)** Full length blots corresponding to KPC689 CD81 and KPC689 Albumin associated with main Figure 5f.

**Supplementary Figure. 4 | Comparison of SE-FPLC yield with gold standard ultracentrifugation technique and quantification of EV recovery rate of SE-FPLC. (A)** Yield calculations based on concentration of particles (particles/mL) n=4 independent biological replicates for MSC cells. **(B)** Yield calculations based on total amount of particles (Total EV amount) n=4 independent biological replicates for MSC cells. Data are presented as mean value ± SD. Statistical analysis was determined by using unpaired two tailed t-test and p=0.0129. Significance was defined as p<0.05 **(C)** Size distribution and concentration (particles/mL) of particles obtained by SE-FPLC (n=3 independent MSC EV collections via SE-FPLC). NTA profiles constructed by the average curve (solid line) and error band (shaded area) represents standard deviation. **(D, E)** The particle mean size and particle mode size in nm of EVs isolated by UC and SE-FPLC (n=4 independent experiments from MSC cell line). Data are presented as mean value ± SD.**(F)** The recovery rate of SE-FPLC was calculated by passing the EV-enriched peak fractions in SE-FPLC column and calculated the recovered extracellular vesicle ratio after SE-FPLC purification to the input EV quantity. (n=3 technical replicates from Panc1 cell line). Data are presented as mean value ± SD. **(G, H)** Representative UV-Vis chromatographs of the recovered EV fractions.

**Supplementary Figure. 5 | Comparison of SE-FPLC yield, Cryo EM analysis of EV morphology isolated using different methods. (A)** Yield calculations based on concentration of particles (particles/mL) across SE-FPLC, SEC and DG isolation methods. n=4 independent biological replicates for Panc1 cells. Data are presented as mean value ± SD. **(B)** Yield calculations based on total amount of particles (Total EV amount) for EVs isolated by SE-FPLC, SEC and DG. n=4 independent biological experiments from Panc1 cells. Data are presented as mean value ± SD. **(C, D)** The particle mean size and particle mode size in nm of EVs isolated by SE-FPLC, SEC and DG (n=4 biological independent experiments from Panc1 cell line). Data are presented as mean value ± SD. **(E)** Validation of SEC isolation method. Western blot analysis of Syntenin-1 and KRAS^G12D^ for Panc1 EVs isolated by SEC Fractions (F7-F20). Fractions F7-F10 were considered EV enriched and had enrichment for EV associated markers Syntenin-1 and KRAS^G12D^ as PDAC EVs. The relative intensity quantification of western blot was analyzed by ImageJ and represented below the western blot. High resolution Cryo EM analysis of Panc1 cell isolated EVs via **(F)** SE-FPLC, **(G)** SEC, **(H)** DG and **(I)** dUC. Representative images are shown in the figures. Scale bars, 200 nm

**Supplementary Figure. 6 | Comparison of total EV proteins by silver staining.(A)** Equal-sample volume (40 μL EV sample) MSC EV-enriched and Pro-Enriched SE- FPLC fractions **(B)** Equal-samples (40 μL EV sample) silver staining analysis of Panc1EV-enriched and Pro-enriched SE-FPLC fractions.

**Supplementary Figure. 7 | Full length unprocessed blot pertaining to Supplementary** Figure 5E**. (A)** Full length blots corresponding to Panc1 Syntenin-1 and **(B)** KRAS^G12D^ associated with supplementary Figure 5e. **(C)** Full length blot corresponding to HPNE KRAS^G12D^ and Syntenin-1 for validating assay specificity. **(D)** Quantification of relative level of Syntenin-1 in blot corresponding to supplementary figure 7C.

**Supplementary Fig. 8 | Cryo-EM imaging and EV size distribution. (A-F)** Cryo-EM images of exosomes isolated from serum samples by SE-FPLC. **(G)** Characterization of EV size distribution based on the Cryo-EM images. 60% of the EV sizes are between 40 and 140 nm. 25% of the EV sizes are between 150nm and 220nm nm. (n=71 independent EVs).

**Supplementary Fig. 9 | Comparison of processing time for different methods for isolation of EVs and SE-FPLC EV enrichment efficiency. (A)** Comparison of processing time of four methods (n=3 independent biological experiments from Panc1 cells). **(B)** Quantification of number of particles in SE-FPLC EV-enriched fraction and SE- FPLC protein-enriched fraction. (n=3 technical replicates for EVs from Panc1 cells).Statistical analysis was determined by using unpaired two-tailed t-test and p<0.0001. Significance was defined as p<0.05.

**Supplementary Fig. 10 | Quantification of protein amounts of EV-enriched peak fractions and protein-enriched peak fractions by SE-FPLC. (A)** Micro-BCA analysis of EV-enriched peak fractions isolated by SE-FPLC. The average protein amount across n=8 measurements was 92.76 μg/mL. **(B)** Micro-BCA analysis of Pro-enriched peak fractions isolated by SE-FPLC. The average protein amount across n=8 measurements was 778.79 μg/mL. **(C)** Ratio of P_ProPeak_/P_EVPeak_ can be used as a quality control measure and the peak ratio had a CV of 16.4% across all measurements. EVs isolated from MSC cell line via SE-FPLC across all measurements.

